# VAMP3 and VAMP8 regulate the development and functionality of parasitophorous vacuoles housing *Leishmania amazonensis*

**DOI:** 10.1101/2020.07.09.195032

**Authors:** Olivier Séguin, Linh Thuy Mai, Sidney W. Whiteheart, Simona Stäger, Albert Descoteaux

## Abstract

To colonize mammalian phagocytic cells, the parasite *Leishmania* remodels phagosomes into parasitophorous vacuoles that can be either tight-fitting individual or communal. The molecular and cellular bases underlying the biogenesis and functionality of these two types of vacuoles are poorly understood. In this study, we investigated the contribution of host cell Soluble N-ethylmaleimide-sensitive-factor Attachment protein REceptor proteins in the expansion and functionality of communal vacuoles as well as on the replication of the parasite. The differential recruitment patterns of Soluble N-ethylmaleimide-sensitive-factor Attachment protein REceptor to communal vacuoles harboring *L. amazonensis* and to individual vacuoles housing *L. major* led us to further investigate the contribution of VAMP3 and VAMP8 in the interaction of *Leishmania* with its host cell. We show that whereas VAMP8 contributes to optimal expansion of communal vacuoles, VAMP3 negatively regulates *L. amazonensis* replication, vacuole size, as well as antigen cross-presentation. In contrast, neither proteins has an impact on the fate of *L. major*. Collectively, our data support a role for both VAMP3 and VAMP8 in the development and functionality of *L. amazonensis*-harboring communal parasitophorous vacuoles.

## INTRODUCTION

*Leishmania* is the protozoan parasite responsible for a spectrum of diseases termed leishmaniasis. Shortly after inoculation into a mammalian host by an infected sand fly, promastigote forms of the parasite are taken up by host phagocytes. To colonize those cells, promastigotes subvert their microbicidal machinery by targeting signaling pathways and altering intracellular trafficking (1–3), and create a hospitable niche that will allow their differentiation and replication as mammalian-stage amastigote forms (4, 5). Most *Leishmania* species replicate in tight-fitting individual parasitophorous vacuoles (PVs), with the exception of species of the *L. mexicana* complex which replicate in spacious communal PVs. For tight-fitting individual PVs, replication of the parasites entails vacuolar expansion and fission, yielding two individual PVs containing one parasite each. In contrast, communal PVs occupy a large volume within infected cells and contain several amastigotes. These two different lifestyles imply that *Leishmania* uses distinct strategies to create the space needed for its replication within infected cells.

Biogenesis and expansion of communal PVs is accomplished through the acquisition of membrane from several intracellular compartments (4). Hence, shortly after phagocytosis, phagosomes harboring *L. amazonensis* fuse extensively with host cell late endosomes/lysosomes and secondary lysosomes (6, 7), consistent with the notion that communal PVs are highly fusogenic (8). Homotypic fusion between *L. amazonensis*-containing PVs also occurs, but its contribution to PV enlargement remains to be further investigated (9, 10). Interaction of these PVs with various sub-cellular compartments indicates that the host cell membrane fusion machinery is central to the biogenesis and expansion of communal PVs and is consistent with a role for Soluble N-ethylmaleimide-sensitive-factor Attachment protein REceptor (SNARE) proteins in this process (11). In this regard, *Leishmania*-harboring communal PVs interact with the host cell endoplasmic reticulum (ER) and disruption of the fusion machinery associated with the ER-Golgi intermediate compartment (ERGIC) was shown to inhibit parasite replication and PV enlargement (12–15). PV size and parasite replication were also shown to be controlled by LYST (16), a protein associated to the integrity of lysosomal size and quantity (17), and by the scavenger receptor CD36, possibly through the modulation of fusion between the PV and endolysosomal vesicles (18). Interestingly, the V-ATPase subunit d2, which does not participate in phagolysosome acidification, was recently shown to control expansion of *L. amazonensis*-harboring PVs through its ability to modulate membrane fusion (19). In addition to regulating the biogenesis of phagolysosomes, trafficking and fusion events play a critical role in the acquisition by phagosomes of the capacity to process antigens for presentation to T cells (20–22). In the case of *Leishmania*-harboring PVs, investigations on their immunological properties revealed that *Leishmania* may interfere with the capacity of these organelles to process antigens for the activation of T cells (23–34).

How parasites of the *L. mexicana* complex co-opt host cell processes to create and maintain hospitable communal PVs is poorly understood. In the case of *Leishmania* species living in tight-fitting individual PVs, two abundant components of the promastigotes surface coat modulate PV composition and properties: the glycolipid lipophosphoglycan (LPG) and the zinc-metalloprotease GP63. LPG contributes to the ability of *L. donovani, L. major*, and *L. infantum* to colonize phagocytes (35–37) by reducing phagosome fusogenecity towards late endosomes and lysosomes, impairing assembly of the NADPH oxidase, and inhibiting phagosome acidification (35, 36, 38–42). In contrast, LPG is not required for infection of macrophages or mice by *L. mexicana* (43) suggesting that LPG has little impact on the formation and properties of communal PV. GP63 contributes to the properties and functionality of tight-fitting PVs by targeting components of the host membrane fusion machinery, including the SNARE proteins VAMP8 and Syt XI, both of which regulate microbicidal and immunological properties of phagosomes (32, 44). We also previously reported that episomal expression of GP63 increases the ability of a *L. mexicana Δcpb* mutant to replicate in macrophages and to generate larger communal PV (45). This correlated with the exclusion of the endocytic SNARE, VAMP3, from the PV, suggesting that this component of the host cell membrane fusion machinery participates in the regulation of communal PVs biogenesis.

In the present study, we investigated the recruitment and trafficking kinetics of host SNAREs to communal PVs induced by *L. amazonensis* and to tight-fitting individual PVs induced by *L. major*. We provide evidence that the endocytic SNAREs VAMP3 and VAMP8 regulate the development and functionality of communal PVs and impact the growth of *L. amazonensis*.

## RESULTS

### Differential recruitment of SNAREs to PVs harboring L. major and L. amazonensis

To study the host cell machinery associated with the development of *Leishmania*-harboring PVs, we first compared the recruitment and trafficking kinetics of host SNAREs to tight-fitting individual and communal PVs. To this end, we infected BMM with either *L. major* strain GLC94 or *L. amazonensis* strain LV79 and used confocal immunofluorescence microscopy to assess the fate of SNAREs associated with various host cell compartments up to 72 h post-infection. As shown in Fig 1A, we found a gradual increase in the proportion of *L. amazonensis* LV79-harboring, communal PVs, positive for the recycling endosomal v-SNARE VAMP3, whereas the proportion of VAMP3-positive tight-fitting PVs harboring *L. major* GLC94 remained below 5%. We observed a similar selective recruitment pattern to communal PVs harboring *L. amazonensis* LV79 for the plasma membrane-associated t-SNARE SNAP23 (Fig 1B), the ER t-SNARE Syntaxin-18 (Fig 1C), and the *trans*-Golgi t-SNARE Vti1a (Fig 1D). The lysosomal-associated protein LAMP1, which we used to define lysosomes (46, 47), also accumulated mainly on communal PVs containing *L. amazonensis* LV79 (Fig. S1A). In contrast, the late endosomal v-SNARE VAMP8 was gradually recruited to tight-fitting PVs containing *L*. *major*, but not to communal PVs containing *L. amazonensis* LV79 (Fig. 1E). In the case of the early endosomal t-SNARE Syntaxin-13, only a small subset (10-15%) of either *L. major* GLC94 or *L. amazonensis* LV79 were housed in a positive PV (Fig S1B). These results support the notion that in contrast to *L. major, L. amazonensis* recruits components from both the host cell endocytic and the secretory pathways for the development and maintenance of communal PVs (4, 13).

**Fig 1.**
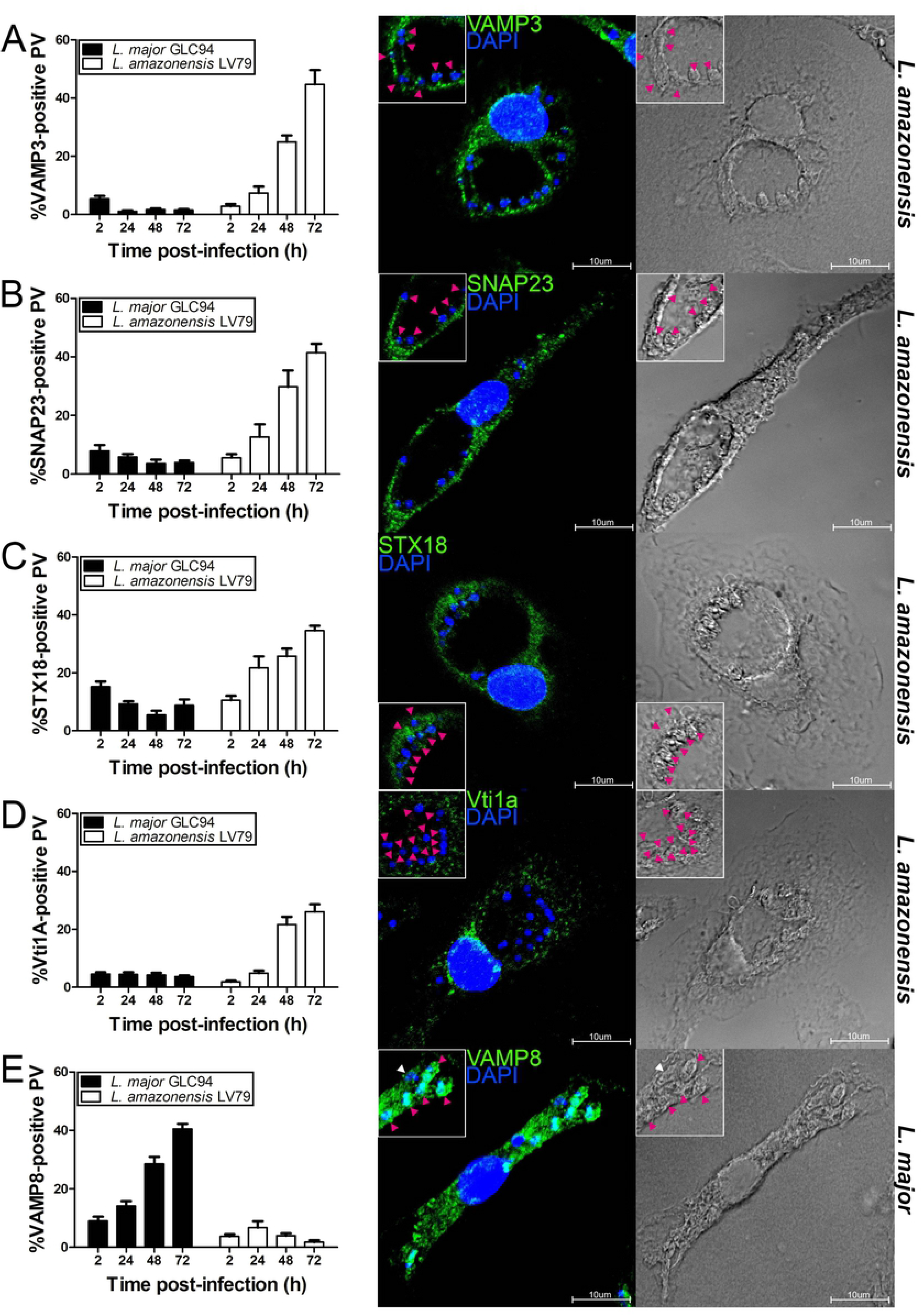
Differential recruitment of SNAREs to *Leishmania*-harboring PVs. BMM were infected with *L. major* GLC94 or *L. amazonensis* LV79 promastigotes and the presence of (A) VAMP3,(B) SNAP23,(C) STX18,(D) Vti1A, and (E) VAMP8 to PVs was assessed and quantified by confocal immunofluoresnce microscopy at 2, 24, 48 and 72 h post-phagocytosis. SNAREs are shown in green and DNA in blue. Data are presented as the mean ± SEM of values from three independent experiments. Representative images of 3 experiments are shown. Insets display PV area. Pink arrowhead indicates recruitment while white arrowheads indicate absence of recruitment. Bar, 10μm.

### VAMP3 and VAMP8 contribute to the control of Leishmania infection

Given the differential recruitment patterns of the endocytic v-SNAREs VAMP3 and VAMP8 to tight-fitting individual and communal PVs, we used them as exemplars to further investigate the development of individual and communal PVs. We first investigated the impact of these two SNAREs on parasite replication. To this end, we infected wild type, *Vamp3^-/-^*, and *Vamp8^-/-^* BMM with either *L. major* GLC94 or *L. amazonensis* LV79 and at various time points, we assessed parasite burden and PV surface area. In the case of VAMP8, we previously reported that its absence had no impact on the replication of *L. major* GLC94 (48). Similarly, absence of VAMP8 had no effect on the replication of *L. amazonensis* LV79 up to 72 h post-infection (Fig 2A). Strikingly, absence of VAMP3 rendered BMM more permissive to the replication of *L. amazonensis* LV79, with nearly twice as many parasites per infected macrophages at 72 h post-infection (Fig 2B). Moreover, PVs harboring *L. amazonensis* LV79 were significantly larger in the absence of VAMP3 (Fig 2C). In contrast, absence of VAMP3 had no impact on the survival and replication of *L. major* GLC94 over a period of 72 h post-infection (Fig 2B). These results indicate a role for VAMP3 in the control of *L. amazonensis* LV79 replication and of communal PVs expansion.

**Fig 2.**
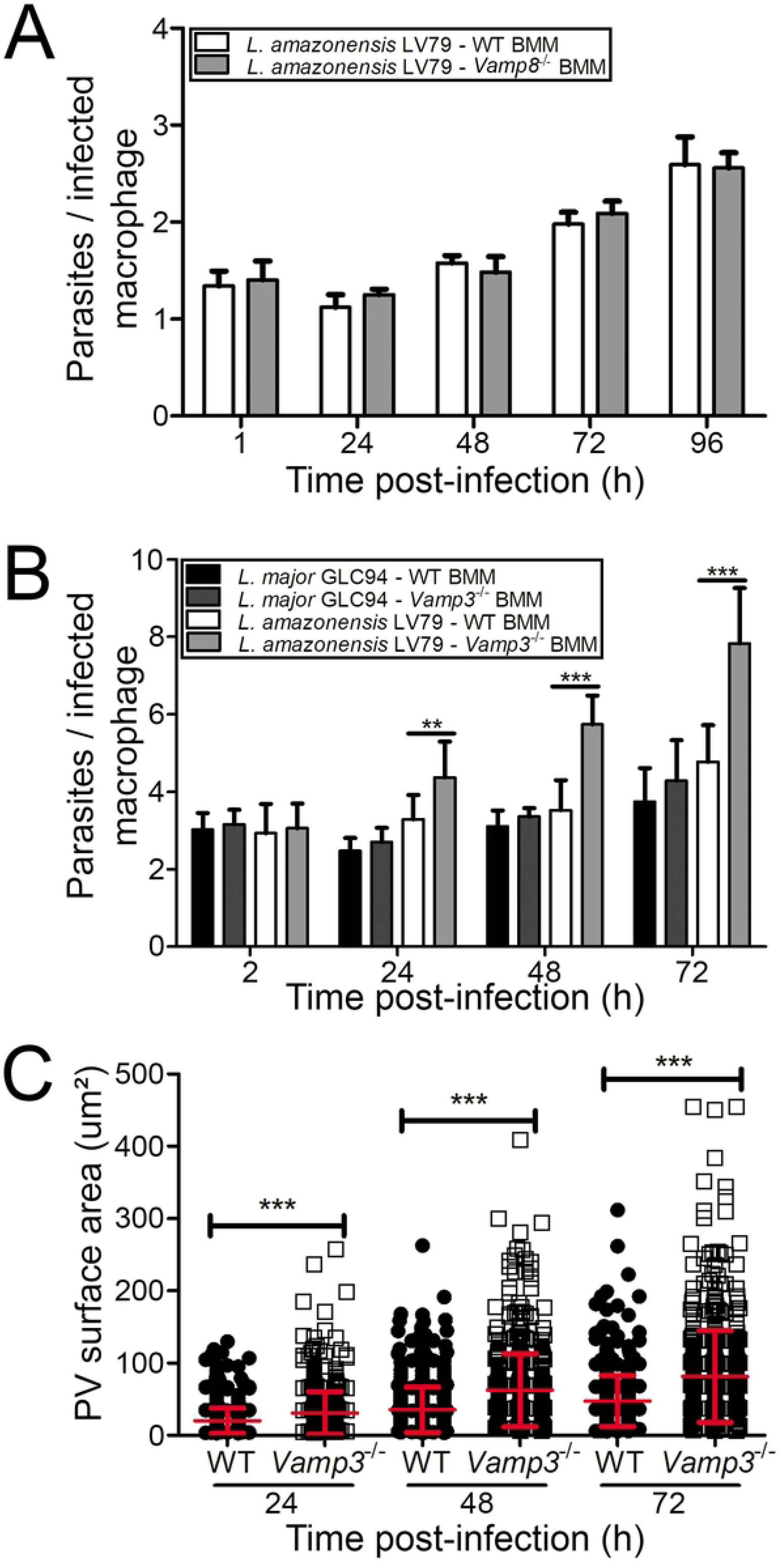
VAMP3 negatively regulates the replication of *L. amazonensis* LV79 and PV expansion. WT, *Vamp8*^-/-^, and *Vamp3*^-/-^ BMM were infected with *L. major* GLC94 or *L. amazonensis* LV79 promastigotes and at various time points post-phagocytosis, parasite replication and PV size were assessed. (A) Quantification of *L. amazonensis* LV79 promastigotes replication in WT or *Vamp8*^-/-^ BMM at 1, 24, 48, 72, and 96 h post-infection. Data are presented as the mean ± SEM of values from three independent experiments. (B) Quantification of *L. major* GLC94 and *L. amazonensis* LV79 replication in WT or *Vamp3*^-/-^ BMM at 2, 24, 48 and 72 h post-infection. Data are presented as the mean ± SEM of values from three independent experiments. **p ≤ 0.01, ***p ≤ 0.001. C. Quantification of PV size in WT and *Vamp3*^-/-^ BMM infected with *L. amazonensis* LV79 at 2, 24, 48 and 72 h post-infection. Data are presented as a cloud with mean ± SD of values from three independent experiments for a total of 450 PV. ***p ≤ 0.001.

A quantitative proteomic analysis has recently been performed on lesion-derived amastigotes of the highly virulent *L. amazonensis* strain PH8 and of strain LV79 (49). Interestingly, the virulence factor GP63 was found to be expressed at higher levels by *L. amazonensis* PH8 amastigotes (49). Our previous report suggesting that GP63 contributes to the expansion of large communal PVs harboring *L. mexicana* (45), led us to compare PVs harboring these two *L. amazonensis* strains. First, we assessed the levels of GP63 expressed by both strains by Western blot analysis of infected BMM lysates for various time points after phagocytosis (Fig 3A). Higher levels of GP63 were present in BMM infected with *L*. amazonensis PH8 compared to BMM infected with *L. amazonensis* LV79 at 2 h post-phagocytosis. In contrast, at later time points (24, 48, and 72 h post-phagocytosis), we detected higher levels of GP63 in BMM infected with LV79 compared to BMM infected with PH8 (Fig. 3A). We observed a similar pattern for the activity of GP63 in lysates of infected BMM (Fig. 3B). Since both VAMP3 and VAMP8 are down-modulated in infected macrophages (32, 45, 48), we assessed the levels of those SNAREs by Western blot analysis of lysates of BMM infected for 2, 24, and 72 h with either strains of *L. amazonensis*. As shown in Fig 3C, both SNAREs were down-modulated by nearly 50% at 72 h post-phagocytosis in infected BMM. Using confocal immunofluorescence microscopy, we compared the kinetics of VAMP3 recruitment to PVs in BMM infected for various time points with either strains of *L. amazonensis*. Remarkably, data shown in Fig 3D revealed that in contrast to PVs harboring *L. amazonensis* LV79, there was no significant recruitment of VAMP3 to PVs containing *L. amazonensis* PH8. Evidence that VAMP3 and VAMP8 may exert overlapping function within the endocytic pathway and that both SNAREs can substitute for each other (50) led us to assess the fate of VAMP8 during infection of BMM with both strains of *L. amazonensis*. Unexpectedly, we observed a significant recruitment of VAMP8 to *L. amazonensis* PH8-harboring PVs at 48 h and 72 h post-infection (Fig 3E). Together, these findings suggest that recruitment of VAMP3 and VAMP8 to *L. amazonensis*-harboring communal PVs is strain-dependent.

**Fig 3.**
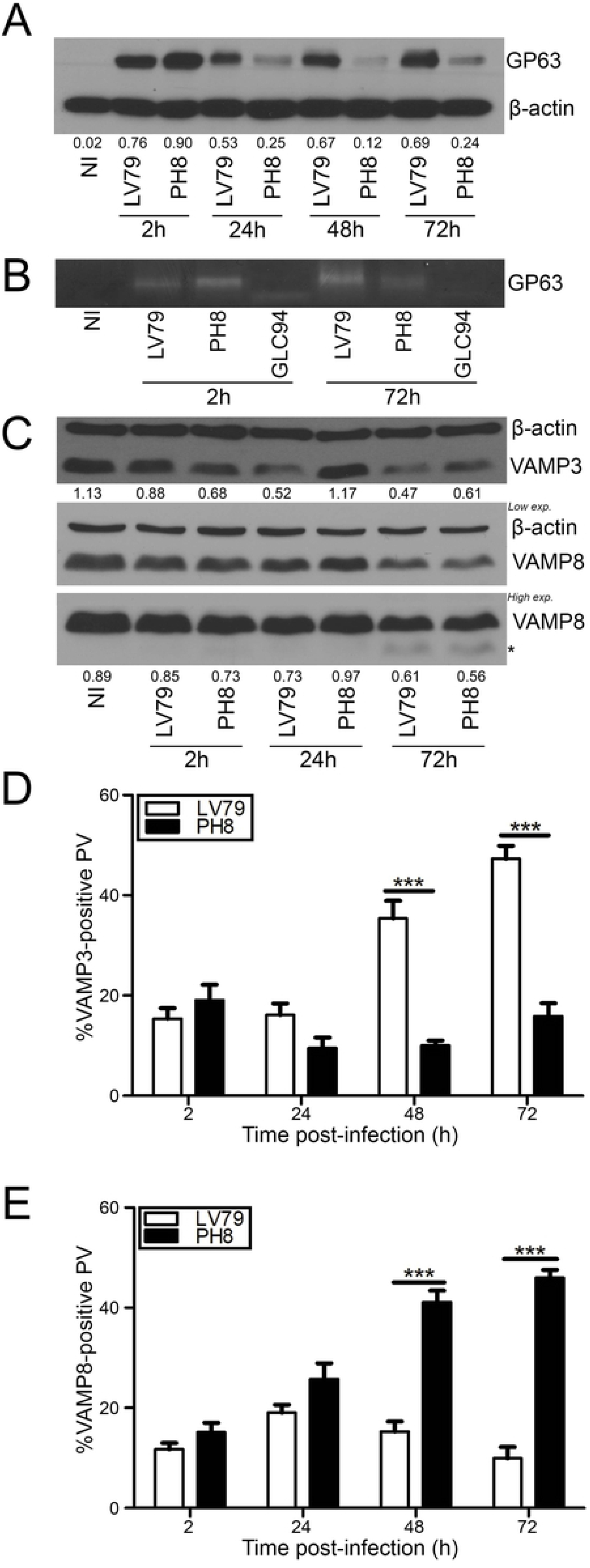
Strain-specific differences in the recruitment of VAMP3 and VAMP8 to PVs harboring *L. amazonensis*. Lysates of BMM infected for 2, 24, 48 and 72 h with *L. amazonensis* LV79 or PH8 promastigotes were prepared. (A) GP63 levels were assessed by western blotting. Representative blot of 3 independent experiments is shown. Band intensities were normalized to β-actin. (B) GP63 activity was determined by zymography. Representative image of 3 experiments is shown. (C) BMM were infected for 2, 24 and 72 h with *L. amazonensis* LV79 or *L. amazonensis* PH8 promastigotes and integrity of both VAMP3 and VAMP8 cleavage was assessed through immunoblot analysis. Representative blots of 3 experiments are shown. Band intensities were normalized to β-actin. *Indicates cleavage fragments. BMM were infected with *L. amazonensis* LV79 or *L. amazonensis* PH8 promastigotes and localization of VAMP3 (D) and VAMP8 (E) to PVs was quantified by confocal microscopy at 2, 24, 48 and 72 h. Data are presented as the mean ± SEM of values from three independent experiments. ***p ≤ 0.001.

Given these strain-dependent differences in the recruitment of VAMP3 and VAMP8 to communal PVs, we assessed the impact of those SNAREs on the replication of both strains of *L. amazonensis* and on PV expansion. We infected either WT, *Vamp3*^-/-^, or *Vamp8*^-/-^ BMM and we determined parasite burden and PV size at various time points post-phagocytosis. As shown in Fig 4A, VAMP3-deficient BMM were more permissive than WT BMM for the replication of both *L. amazonensis* strains. Additionally, for both strains, absence of VAMP3 led to the formation of larger PVs at 48 h and 72 h post-infection (Fig 4B), consistent with a role for that SNARE in the control of PV size and *L. amazonensis* replication. In contrast, whereas absence of VAMP8 had no impact on the replication of *L. amazonensis* LV79, it significantly restricted the replication of strain PH8 at 24, 48, and 72 h post-phagocytosis (Fig 5A). These results suggest that VAMP8 participates in fusion events required for the replication of *L. amazonensis* PH8. Interestingly, for both *L. amazonensis* strains, PV size was reduced in the absence of VAMP8 (Fig 5B), indicating a role for this SNARE in the regulation of PV expansion. In addition, these results suggest that there is no direct correlation between PV size and parasite replication.

**Fig 4.**
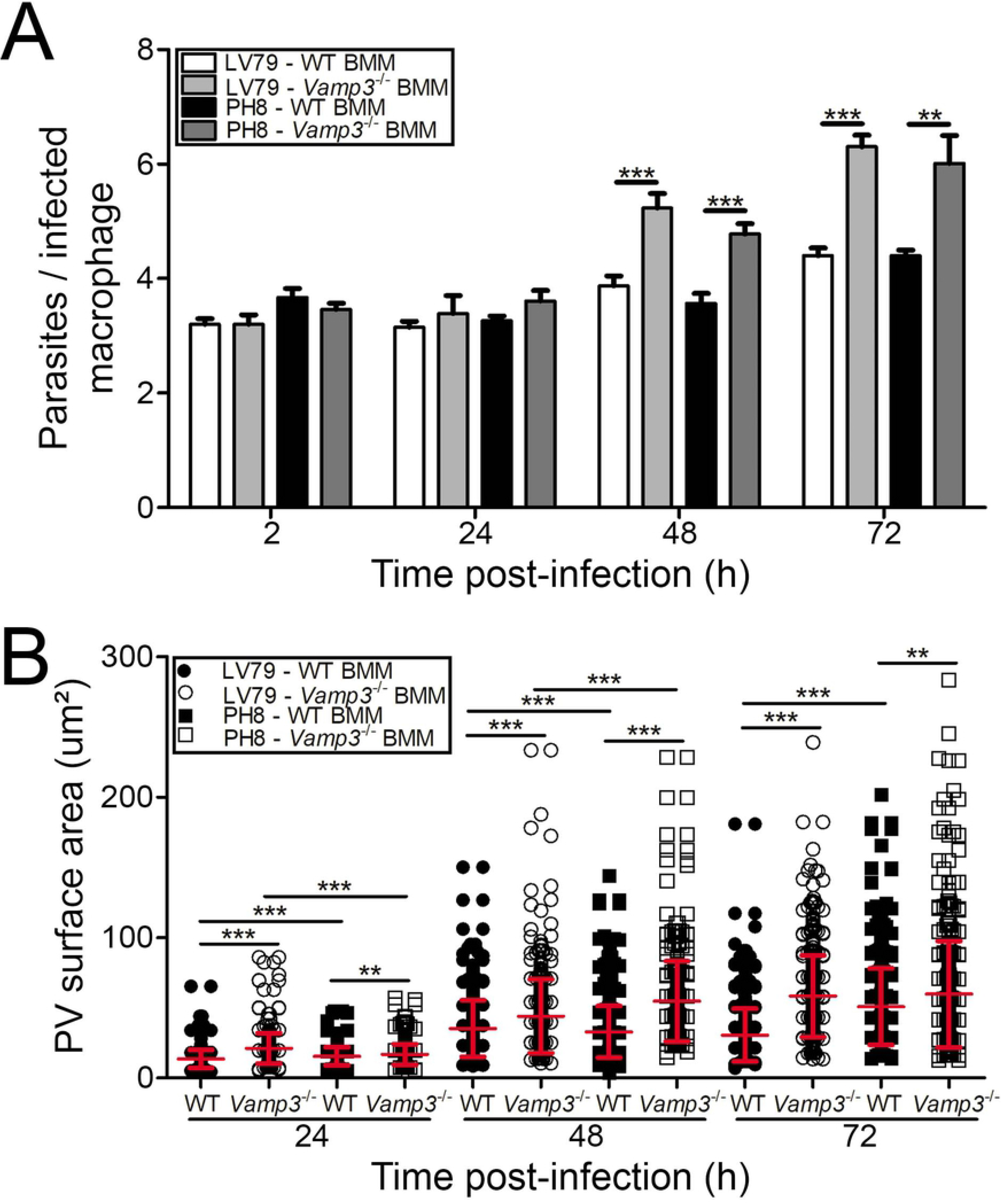
Impact of VAMP3 on the replication of *L. amazonensis* LV79 and PH8 and PV size. WT and *Vamp3*^-/-^ BMM were infected with *L. amazonensis* LV79 and PH8 promastigotes and at various time points post-phagocytosis, parasite replication and PV size were assessed. (A) Parasite replication, data presented as the mean ± SEM of values from three independent experiments. **p ≤ 0.01, ***p ≤ 0.001. (B) PV surface area, data are presented as a cloud with mean ± SD of values from three independent experiments for a total of 450 PV. **p ≤ 0.01, ***p ≤ 0.001.

**Fig 5.**
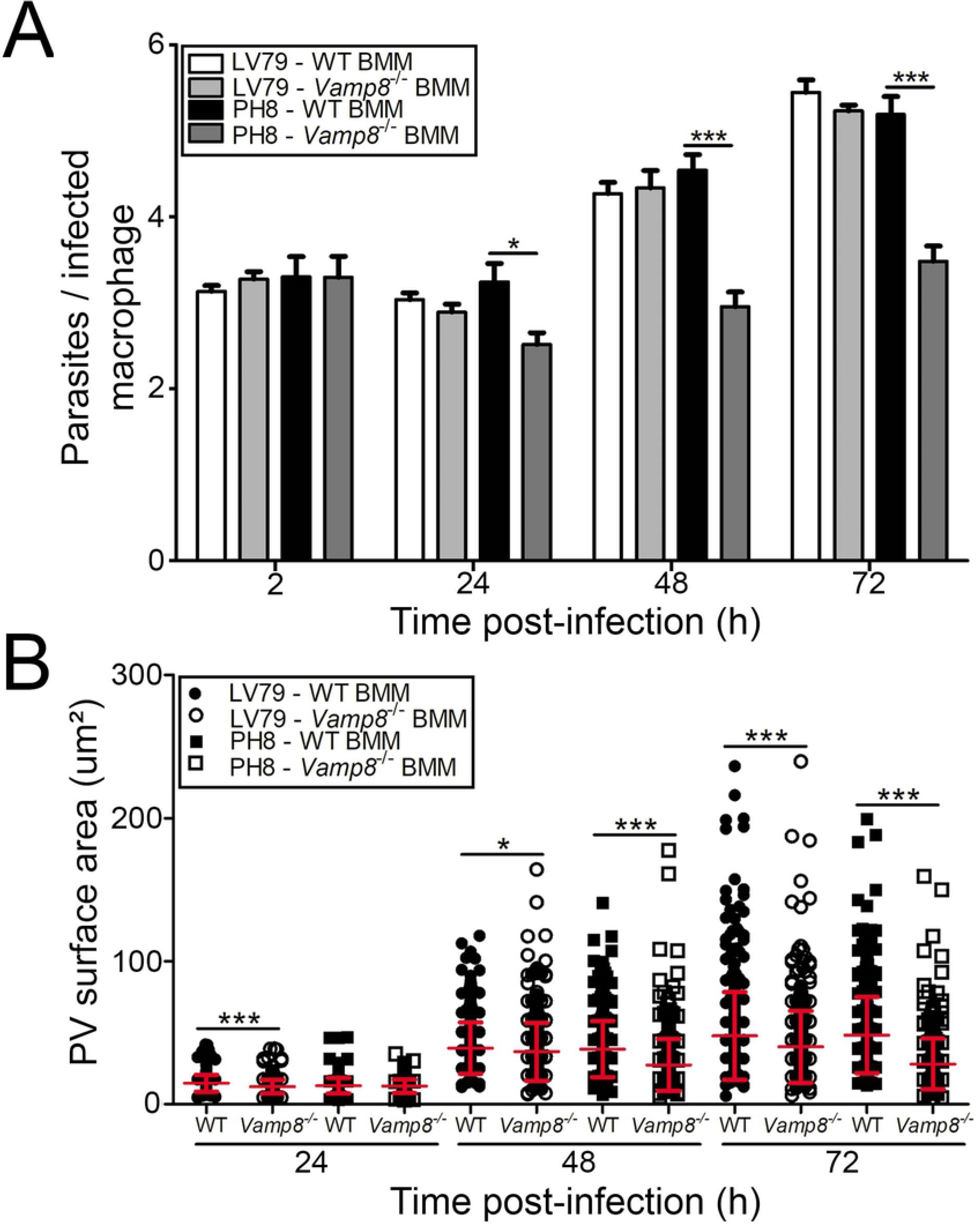
Impact of VAMP8 on the replication of *L. amazonensis* LV79 and PH8 and PV size. WT and *Vamp8^-/-^* BMM were infected with *L. amazonensis* LV79 and PH8 promastigotes and at various time points post-phagocytosis, parasite replication and PV size were assessed. (A) Parasite replication, data presented as the mean ± SEM of values from three independent experiments. *p ≤ 0.05, ***p ≤ 0.001. (B) PV surface area, data are presented as a cloud with mean ± SD of values from three independent experiments for a total of 450 PV. *p ≤ 0.05, ***p ≤ 0.001.

### VAMP3 negatively regulates antigen cross-presentation

Given the impact of VAMP3 on parasite replication and PV size, we sought to investigate whether absence of VAMP3 alters communal PV functionality. We therefore elected to focus on antigen cross-presentation, a phagosomal function which has received little attention in the context of *L. amazonensis* infection. To this end, we first compared the capacity of WT and *Vamp3*^-/-^bone marrow-derived dendritic cells (BMDC) to cross-present ovalbumin following internalization of OVA-coated latex beads. As shown in Fig 6A, at 6 h post-phagocytosis, OVA-pulsed WT and *Vamp3^-/-^* BMDC were equally efficient at activating OVA_257-264_-specific B3Z hybridomas, as previously reported for BMDC fed with *E. coli*-OVA (51). Next, we generated *L. major* GLC94 and *L. amazonensis* strains LV79 and PH8 expressing ovalbumin (OVA) at similar levels (Fig 6B) and used them to infect WT and *Vamp3^-/-^* BMDC for 48 h prior to adding OVA-specific OT-I CD8 T cells (52). At this time, both *L. amazonensis*-OVA strains were present in enlarged PVs, whereas *L. major*-OVA replicated in tight individual PVs (Fig S2). We then assessed cross-presentation by measuring the expression of T cell activation markers CD69 and CD44 on OT-I T cells following exposure to *L. amazonensis*-OVA- and *L.major*-OVA-infected BMDC. As shown in Figures 6C-E, *L. major*-OVA and *L. amazonensis*-OVA efficiently suppressed the ability of WT infected BMDC to cross-present ovalbumin and activate OT-I CD8 T cells. Interestingly, significantly lower frequencies of CD44^+^ OT-I T cells (Fig. 6C) and CD69 MFI levels (Fig. 6D) were observed for OT-I T cells exposed to *L. amazonensis*-OVA-infected compared to *L. major*-infected BMDC, suggesting that *L. amazonensis* may be more efficient at inhibiting cross-presentation than *L. major*. Similar to WT BMDC, *L. major*-OVA-infected *Vamp3^-/-^* BMDC also failed to prime OT-I T cells. In contrast, cross-presentation was significantly increased in VAMP3-deficient BMDC infected with either strain of *L. amazonensis*-OVA. Collectively, our results suggest that *L. amazonensis* efficiently suppresses antigen cross-presentation in BMDC, and that VAMP3 negatively regulates the ability of communal PVs harboring *L. amazonensis* to cross-present antigens.

**Fig 6.**
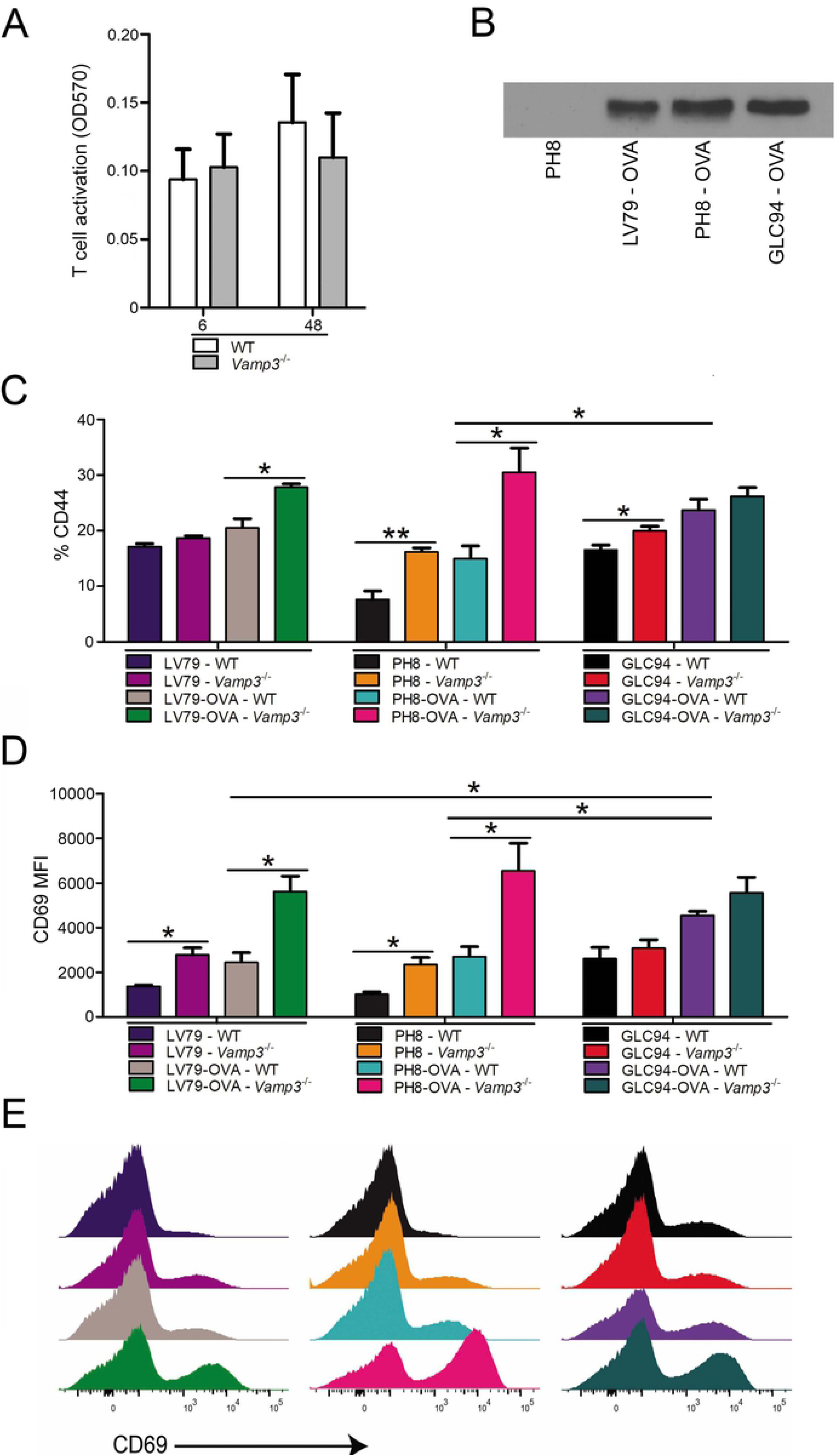
VAMP3 negatively regulates antigen cross-presentation from communal PVs harboring *L. amazonensis*. (A) WT and *Vamp3^-/-^* BMDC were fed OVA-coated beads and cross-presentation was evaluated using B3Z T cell hybridoma. Data are presented as the mean ± SEM of values from three independent experiments. (B) Expression of ovalbumin by *L. amazonensis*-OVA LV79, *L. amazonensis*-OVA PH8 and *L. major*-OVA GLC94 was assessed by western blotting. Representative blot of two experiments is shown. (C) and (D) WT and *Vamp3^-/-^* BMDC were infected with either *L. major* GLC94, *L. amazonensis* LV79 and *L. amazonensis* PH8 or *L. major*-OVA GLC94, *L. amazonensis*-OVA LV79, and *L. amazonensis*-OVA PH8 for 48 h. Infected and control uninfected BMDC were incubated with OT-1 T cells and cross-presentation was assessed by measuring expression of the activation markers CD44 (C) and CD69 (D, E) on OT-I T cells by flow cytometry. Graphs show one representative experiment of 6 independent experiments *p ≤ 0.05, **p ≤ 0.01.

## DISCUSSION

Biogenesis and functionality of phagosomes depend on intracellular vesicular trafficking and membrane fusion events mediated by SNAREs (11, 32, 53, 54). In this study, we compared and analyzed the recruitment and trafficking kinetics of host cell SNAREs to tight-fitting individual PVs induced by *L. major* and to large communal PVs induced by *L. amazonensis*. Our results revealed differences in the components of host cell membrane fusion machinery associated to these two types of PVs, consistent with the notion that tight-fitting individual PVs and large communal PVs differ in their capacity to interact with host cell compartments. Moreover, we obtained evidence that both VAMP3 and VAMP8 regulate the development and functionality of *L. amazonensis*-harboring PVs.

Previous studies revealed that although VAMP3 plays no major role in phagocytosis (55), this SNARE contributes to host defense against infections by regulating the delivery of TNF at the nascent phagocytic cup through focal exocytosis and by contributing to the formation of autophagosomes during xenophagy (56–58). Moreover, VAMP3 may be co-opted by vacuolar pathogens to create and develop their intracellular replicative niches. Hence, in macrophages infected with *Yersinia pseudotuberculosis*, VAMP3 participates in the formation of single-membrane LC3-positive vacuoles containing the bacteria, although it is not known whether VAMP3 influences *Yersinia* replication (59). Using host cells co-expressing EGFP-VAMP3 and tetanus toxin (which cleaves VAMP3), Campoy and colleagues obtained evidence that VAMP3 is involved in the biogenesis and enlargement of the *Coxiella burnetti* replicative vacuoles, by mediating fusion of these replicative vacuoles with multivesicular bodies (60). In the case of *Brucella melitensis*, although infection increases VAMP3 expression in the mouse macrophage cell line J774, its silencing had no effect on the survival and replication of the bacteria, indicating that VAMP3 is not essential for the biogenesis and expansion of *Brucella*-containing vacuoles (61). For *Leishmania*, our results revealed no role for VAMP3 in the replication of *L. major*, which resides in individual tight-fitting PVs. In contrast, we found that VAMP3 is gradually recruited to *L. amazonensis* LV79-harboring PV starting at 24 h post-infection, when PV expansion becomes noticeable. Unexpectedly, we observed increased replication of *L. amazonensis* and increased PV size in the absence of VAMP3, suggesting that this SNARE may be part of a complex that controls the development of communal PVs, possibly by negatively regulating membrane fusion events as we previously reported for the regulator of membrane fusion complexes synaptotagmin XI (62). Of interest, a negative regulatory role for VAMP3 has recently been reported in platelets, where absence of VAMP3 led to enhanced platelet spreading and clot retraction (63). In this context, it was proposed that absence of VAMP-3 could increase the formation of other SNARE complexes and thus increase the exocytosis events needed for spreading. One may thus envision that absence of VAMP3 may favor the formation of SNARE complexes that lead to increased fusion events required for PV expansion. It is also possible that VAMP3 acts as an inhibitory SNARE (or i-SNARE) (64), where it would compete with and substitute for a fusogenic subunit, thereby inhibiting fusion and limiting PV expansion. Alternatively, VAMP3 may mediate the delivery of an anti-microbial cargo to PV harboring *L. amazonensis*, thereby limiting parasite replication and PV expansion. Clearly, our study revealed a novel role for VAMP3 in the control of communal PV expansion and *L. amazonensis* replication, highlighting the fact that depending of the pathogen, a molecule involved in regulating vesicular trafficking and membrane fusion events may either favor or impair pathogen growth, as recently described for Rab11 in the context of *L. pneumophila* infection (65). Further studies will be aimed at elucidating the underlying mechanism(s).

Our previous work revealed that whereas VAMP8 has no influence on the survival and replication of *L. major*, it contributes to the ability of phagosomes to cross-present antigens by regulating the phagosomal recruitment of NOX2 (32, 48). Here, we obtained evidence that absence of VAMP8 leads to reduced expansion of communal PVs harboring *L. amazonensis*. This suggests that VAMP8 participates to the recruitment of membrane required for the expansion of communal PVs by mediating their interactions with endosomes/lysosomes, and/or by mediating homotypic fusion among communal PVs. The observation that reduction of PV size in the absence of VAMP8 impacted the replication *L. amazonensis* PH8, but not of *L. amazonensis* LV79, is intriguing and suggests that *L amazonensis* replication does not absolutely correlate with PV expansion. Hence, whether parasite growth is the main signal governing PV expansion remains a lingering question (27) for which there has been little data to provide a clear answer. It will thus be of interest to further investigate the nature of the host factors and parasite effectors that modulate PV expansion. The different fates of *L. amazonensis* LV79 and PH8 in *Vamp8^-/-^* macrophages illustrate the perils of drawing conclusions based on experiments performed with a single *Leishmania* strain or isolate (66).

Cross-presentation of microbial peptides on major histocompatibility (MHC) I molecules is an important host defense mechanism aimed at deploying CD8^+^ T cells responses against intracellular pathogens, including *Leishmania* (67–73). Previous studies revealed that phagosomes acquire, through a series of interactions with other organelles, the machinery required to become self-sufficient for antigen cross-presentation (20–22). However, the involvement of the various vesicular trafficking pathways in this process remains to be fully understood (72, 74). To date, a number of SNAREs associated to these pathways have been shown to regulate the trafficking events involved in the acquisition of the phagosomal cross-presentation machinery, including the ER/ERGIC SNARE Sec22b (54) and the endocytic SNAREs VAMP8 and SNAP-23 (32, 51). Sec22b was shown to mediate the delivery of the MHC-I peptide loading complex from the ERGIC to phagosomes through pairing with the plasma membrane SNARE syntaxin-4 present on phagosomes (54). Recruitment of the NOX2, whose activity is crucial to regulate the levels of phagosomal proteolysis required for optimal cross-presentation (75), is mediated by both VAMP8 and SNAP-23 (32, 76). Our finding that absence of VAMP3 increases the level of cross-presentation by *L. amazonensis*-harboring communal PVs illustrates the complex regulation of this process and indicates that communal PVs do not behave like model phagosomes used to define the molecular bases of cross-presentation (20, 22, 77). Further characterization of communal PVs induced by *L. amazonensis* and of the role of VAMP3 in their formation may therefore yield novel information on the process of antigen cross-presentation in the context of cells infected with a pathogen residing in communal vacuoles.

We provided evidence that both VAMP3 and VAMP8 participates in the development and functionality of *L. amazonensis*-harboring communal PVs. Whereas VAMP3 has a detrimental impact on parasite replication, PV size, and antigen cross-presentation, VAMP8 contributes to PV expansion but does not affect replication. In both cases, the exact mechanisms remain to be elucidated. This is an interesting issue since both VAMP3 and VAMP8 have previously been shown to exert overlapping function and they can substitute for each other (50). Depending on the cell type and of the intracellular compartment, both SNAREs form complexes with SNAP-23 and various syntaxins. To shed more light on the biology of *L. amazonensis*-harboring PVs, future studies will be aimed at identifying the *trans*-SNARE complexes formed by VAMP3 and VAMP8 during biogenesis and expansion of communal PVs. Since phagosomes play a central role in innate and adaptive immunity, a better understanding of the biology of communal PVs containing *L. amazonensis* may provide new insights into the mechanisms used by this parasite to develop in a communal PV and to evade the immune system, which may be useful for the design of future interventions to prevent or treat infection.

## MATERIALS AND METHODS

### Ethics statement

Experiments involving mice were done as prescribed by protocol 1406-02, which was approved by the *Comité Institutionnel de Protection des Animaux* of the Institut national de la recherche scientifique. These protocols respect procedures on good animal practice provided by the Canadian Council on animal care. *Vamp8*^-/-^ mice were obtained from Dr Wan Jin Hong (A Star Institute, Singapore), *Vamp3^-/-^* mice were obtained from Dr Sidney Whiteheart (University of Kentucky), and OT-I mice were purchased from Jackson Laboratory. All mice were bred and housed at the Institut National de la Recherche Scientifique (INRS) animal facility under specific pathogen-free conditions and used at 6-12 weeks of age.

### Antibodies

The rabbit polyclonal anti-VAMP3, anti-VAMP8, anti-SNAP23, anti-Syntaxin 13, anti-Syntaxin 18 and the guinea pig polyclonal anti-Vti1b antibodies were obtained from Synaptic Systems (SySy). The rat monoclonal anti-LAMP-1 antibody was developed by J. T. August (1D4B) and obtained through the Developmental Studies Hybridoma Bank at the University of Iowa, and the National Institute of Child Health and Human Development. The mouse monoclonal anti-GP63 antibody was provided by Dr. W. R. McMaster (University of British Columbia). FACS analysis were completed with fluorochome conjugated antibodies against CD3-PECy7 (clone 14S-2C11; BD Bioscience), CD8-PB (clone 53-6.7; BD Bioscience), CD44-APC (clone IM7; BD Bioscience) and CD69-PE (clone H1.2F3; eBioscience).

### Bone marrow-derived macrophages and dendritic cells

We used *Vamp8^-/-^* (50) and *Vamp3^-/-^* (78) mice which were maintained on a mixed C57BL/6-129/Sv/J background, and wild type mice were matched littermates. Bone marrow-derived macrophages (BMM) were differentiated from the bone marrow of 6- to 8-week-old mice. Cells were differentiated in complete medium (DMEM [Life Technologies] supplemented with L-glutamine [Life Technologies], 10% heat-inactivated FBS [Gibco], 10 mM HEPES [Bioshop] at pH 7.4, and antibiotics [Life Technologies]) containing 15% v/v L929 cell–conditioned medium as a source of M-CSF at 37°C in a humidified incubator with 5% CO_2_ for a week. To render BMM quiescent prior to experiments, cells were transferred to 6- or 24-well tissue culture microplates (TrueLine) and kept for 16 h in complete DMEM without L929 cell-conditioned medium. Bone marrow-derived dendritic cells (BMDC) were differentiated from the bone marrow of 6- to 8-week-old mice. Cells were differentiated in complete medium (RPMI [Life Technologies] supplemented with 10% heat-inactivated FBS, 10 mM HEPES [Bioshop] at pH7.4 and antibiotics [Life Technologies]) containing 10% v/v GM-CSF at 37°C in a humidified incubator with 5% CO_2_ for a week. Sixteen hours prior to infection, non-adherent cells were transferred in 96-well tissue culture microplates (TrueLine) and kept in RPMI-1640 containing 10% heat-inactivated FBS and 5% v/v GM-CSF.

### Parasite strains and culture

The *Leishmania* strains used in this study were *L. major* GLC94 (MHOM/TN/95/GLC94 zymodeme MON25, obtained from Dr Guizani-Tabbane, Institut Pasteur de Tunis), *L*. amazonensis LV79 (MPRO/BR/72/M1841, obtained from the American Type Culture Collection), and *L. amazonensis* PH8 (IFLA/BR/67/PH8, obtained from the American Type Culture Collection). Promastigotes were obtained from lesion-derived amastigotes and were cultured in *Leishmania* medium (Medium 199 [Sigma-Aldrich] with 10% heat-inactivated FBS, 40 mM HEPES at pH 7.4, 100 μM hypoxanthine, 5 μM hemin, 3 μM biopterin, 1 μM biotin, and antibiotics), in an incubator at 26°C. Promastigotes expressing a secreted form of OVA (*L. major*-OVA and *L. amazonensis*-OVA) were generated by electroporating the pKS-NEO SP:OVA construct, which encodes a fusion protein containing the signal peptide of the *L. donovani* 3′ nucleotidase-nuclease fused to a portion of OVA protein (139–386) containing both MHC class I OVA257-264- and class II OVA323-339-restricted epitopes (79)kindly provided by Dr Alain Debrabant, FDA). Transfected parasites were grown in *Leishmania* medium supplemented with 50 μg/ml G418.

### Infection of macrophages

Promastigotes in late stationary phase were opsonised with C5-deficient serum from DBA/2 mice prior to infections. Phagocytosis was synchronized by incubating macrophages and parasites at 4°C for 10 min and spun at 167 *g* for 1 min. Internalization was then triggered by transferring the plates to 34°C. Two hours post-infection, macrophages were washed twice with complete DMEM to remove non-internalized parasites. Cells were then prepared for confocal immunofluorescence microscopy.

### Confocal immunofluorescence microscopy

Cells adhered to coverslips were fixed with 2% paraformaldehyde (Canemco and Mirvac) for 40 min and blocked/permeabilized for 17 min with a solution of 0.05% saponin, 1% BSA, 6% skim milk, 2% goat serum, and 50% FBS followed by 2 h incubation with primary antibodies and a subsequent incubation with a suitable secondary antibodies in PBS for 45 min (anti-rabbit Alexa Fluor 488 and anti-rat 568; Molecular Probes) and DAPI in PBS for 15 min (Life technologies). Three washes in PBS took place after every step. After the final washes, Fluoromount-G (Southern Biotechnology Associates) was used to mount coverslips on glass slides, and coverslips were sealed with nail polish (Sally Hansen). Macrophages were visualised with the LSM780 microscope 63X objective (Carl Zeiss Microimaging) and images were taken in sequential scanning mode. Image analysis and vacuole size measurements were performed with the ZEN 2012 software. The vacuole size measurements were accomplished via the closed Bezier tool on ZEN 2012 that calculates the surface of a selected area on LAMP-1 and DAPI stained coverslips.

### Lysis, SDS-PAGE and Western blotting

Adherent macrophages in 6-well plates were washed with PBS containing 1 mM sodium orthovanadate and 10 mM 1,10-phenanthroline (Roche) on ice prior to lysis. Cells were then scraped in lysis buffer containing 1% Nonidet P-40 (Caledon), 50 mM Tris-HCl (pH 7.5) (Bioshop), 150 mM NaCl, 1 mM EDTA (pH 8), 10 mM 1,10-phenanthroline, and phosphatase and protease inhibitors (Roche). The lysates were left on ice for 10 min then stored at −70°C. Lysates were thawed on ice and centrifuged for 10 min to remove insoluble matter and then quantified. 10ug of samples were boiled (100°C) for 6 min in SDS sample buffer and migrated in 10% SDS-PAGE gels then transferred onto Hybond-ECL membranes (Amersham Biosciences). The membrane was subsequently blocked for 1h in TBS1X-0.1% Tween containing 5% skim milk, incubated overnight at 4°C with primary antibodies (diluted in TBS 1X-0.1% Tween containing 5% BSA) and incubated 1 h at room temperature with appropriate HRP-conjugated secondary antibodies. 3 washes took place after every step. Membranes were finally incubated in ECL (GE Healthcare) and immunodetection was achieved via chemiluminescence.

### Antigen cross-presentation

BMDC were infected for 48 h with WT *L. amazonensis* LV79-OVA, *L. amazonensis* PH8-OVA or *L. major* GLC94-OVA promastigotes, or with promastigotes not expressing OVA. Cells were then washed and fixed for 5 min at 23°C with 1% (w/v) paraformaldehyde, followed by three washes in complete medium containing 0.1 M glycine. OT-I T cells were enriched from splenocytes of OT-I mice by magnetic cell sorting (MACS) using a CD8^+^ T cell isolation kit (Miltenyi Biotech) as previously described (80). They were then added to the culture for 16 h. The SIINFEKL peptide was used as control for T cell activation and expansion. Antigen cross-presentation was assessed by measuring surface expression modulation of CD69 and CD44 within the CD3^+^CD8α^+^Vα2^+^ population as markers for T cell activation. Cells were analysed after fixation with 2% (w/v) paraformaldehyde using the BD LSR Fortessa flow cytometer (Becton Dickinson). Samples were analyzed with Flowjo software.

For OVA latex bead assays, at 6 h post-infection, 10^5^ BMDC were incubated with uncoated or OVA-coated 0.8 μm latex beads for 1 h of pulse. Cells were incubated for a 3 h chase, fixed with 1% (w/v) paraformaldehyde, and washed in complete medium containing 0.1 M glycine. Cells were then cultured for 16 h at 37°C together with 10^5^ B3Z cells for analysis of T cell activation. B3Z cells, which express β-galactosidase upon specific recognition of the OVA_257-264_ (SIINFEKL)-H-2K^b^ complex, were washed in PBS and lysed (0.125 M Tris base, 0.01 M cyclohexane diaminotetraacetic acid, 50% [v/v] glycerol, 0.025% [v/v] Triton X-100, and 3 mM dithiothreitol [pH 7.8]). A β-galactosidase substrate buffer (1 mM MgSO_4_× 7 H_2_O, 10 mM KCl, 0.39 M NaH_2_PO_4_xH_2_O, 0.6 M Na_2_HPO_4_ ×7 H_2_O, 100 mM 2-mercaptoethanol, and 0.15 mM CPRG [pH 7.8]) was added for 2-4 h at 37°C. Cleavage of CPRG was quantified in a spectrophotometer as absorbance at 570 nm, reflecting T cell activation after cross-presentation.

## Acknowledgements

We thank W. J. Hong for the *Vamp8^-/-^* mice, L. Guizani-Tabbane for the *L. major* GLC94 strain, W.R. McMaster for the anti-GP63 antibody, and J. Tremblay for assistance in immunofluorescence experiments.

## Supporting information captions

**Fig S1. Recruitment of LAMP1 and Syntaxin-13 to PVs**

BMM were infected with serum-opsonized *L. major* GLC94 or *L. amazonensis* LV79 promastigotes and the presence of (A) LAMP1 and (B) Stx13 to PVs was assessed and quantified by confocal microscopy at 2, 24, 48 and 72 h post-phagocytosis. LAMP1 is shown in red, STX13 in green and DNA in blue. Data are presented as the mean ± SEM of values from three independent experiments. Representative images of 3 experiments are shown. Insets display PV area. Pink arrowhead indicates recruitment while white arrowheads indicate absence of recruitment. Bar, 10μm.

**Fig S2. OVA-expressing *Leishmania* replicate in BMDC**

Giemsa-stained WT or *Vamp3^-/-^* BMDC infected for 48 h with serum-opsonized OVA-expressing *L. amazonensis* LV79, *L. amazonensis* PH8, and *L. major* GLC9.

